# Adduction Induces Large Optic Nerve Head Deformations in Subjects with Normal Tension Glaucoma

**DOI:** 10.1101/2021.08.25.457300

**Authors:** Thanadet Chuangsuwanich, Tin A. Tun, Fabian A. Braeu, Xiaofei Wang, Zhi Yun Chin, Satish K. Panda, Martin Buist, Dan Milea, Nicholas Strouthidis, Shamira A. Perera, Monisha E. Nongpiur, Tin Aung, Michael JA Girard

## Abstract

Purpose: To assess optic nerve head (ONH) deformations and strains during adduction, abduction, and intraocular pressure (IOP) elevation in subjects with high-tension glaucoma (HTG) and normal-tension glaucoma (NTG). Design: Clinic-based cross-sectional study. Participants: 114 HTG subjects and 114 NTG subjects. Methods. We recruited 228 subjects (114 subjects with HTG [pre-treatment IOP > 21mmHg] and 114 with NTG [pre-treatment IOP < 21mmHg]). For each subject, we imaged the ONH using spectral-domain optical coherence tomography (OCT) under the following conditions: (1) primary gaze, (2) 20 degree adduction, (3) 20 degree abduction, and (4) primary gaze with acute IOP elevation (to approximately 33 mmHg) achieved through ophthalmodynamometry. For each OCT volume, we automatically segmented the prelaminar tissue (PLT), the choroid, the sclera and the lamina cribrosa (LC) using a deep learning algorithm. We also digitally aligned the OCT volumes obtained from (2)-(4) to the primary gaze volume (1) before performing digital volume correlation (DVC) analysis to quantify IOP- and gaze-induced ONH tissues three-dimensional displacements and effective strain (a local measure of tissue deformation) for all scenarios. Main Outcome Measures: Three-dimensional ONH displacements and strains. Results: Across all subjects, adduction generated high effective strain (4.2 ± 1.4%) in the ONH tissues with no significant difference (p>0.05) with those induced by IOP elevation (4.5 ± 1.5%); while abduction generated significantly lower (p = 0.014) effective strain (3.8 ± 1.1%). Interestingly, the LC of HTG subjects exhibited significantly higher effective strain than those of NTG subjects under IOP elevation (HTG:4.6 ± 1.7% vs NTG:4.1 ± 1.5%, p = 0.047). Conversely, the LC tissue of NTG subjects exhibited significantly higher effective strain than those of HTG subjects under adduction (NTG: 4.9 ± 1.9% vs HTG: 4.0 ± 1.4%, p = 0.041). Conclusion: We found that adduction produced comparable strains and displacements as IOP elevation. We also found that NTG subjects experienced higher strains due to adduction than HTG subjects, while HTG subjects experienced higher strain due to IOP elevation than NTG subjects - and that these differences were most pronounced in the LC tissue.

## Introduction

The standard biomechanical theory of glaucoma hypothesizes that biomechanical forces induced by intraocular pressure (IOP) and cerebrospinal fluid pressure (CSFP) deform the optic nerve head (ONH) tissues, especially at the level of the lamina cribrosa (LC), yielding retinal ganglion cell (RGC) death.^1^ However, IOP and CSFP are not the only loads that can significantly deform the ONH. Biomechanical forces exerted by extraocular muscles during eye movements have been shown to induce significant ONH deformations.^2^ Wang et al. have quantified the effective strain in the LC during eye movements *in vivo* and reported that adduction could induce as much effective strain (i.e. deformation) to the ONH tissue as would an IOP elevation to 40 mmHg.^3^ This is because the optic nerve can become ‘taut’ during adduction and exert a significant traction force to the ONH tissues, as was evidenced through MRI and finite element studies.^4–7^ The functional consequences of such a force are yet unknown.

With the high prevalence of normal tension glaucoma (NTG), especially in some Asian populations^8,9^, the IOP-centric biomechanical theory of glaucoma is insufficient to explain the disease etiology. Vascular deficiency in NTG patients has been proposed as a potential contributing factor,^10^ but its evidence is still not conclusive^11^. From a biomechanical perspective, a few other IOP-independent factors could contribute to the development of NTG; for instance, a low CSFP^12, 13^, structural weaknesses of ocular tissues^14, 15^,an increased susceptibility to optic nerve traction during eye movements^3,5^, or a combination of the aforementioned factors. To date, no studies have compared the biomechanical effects of optic nerve traction in NTG and high-tension glaucoma (HTG) subjects in a relatively large cohort. With increasing evidence that eye movements could induce significant deformation in the ONH, both observed *in vivo*^*3*^ and via computational modelling^16^, we believe that such a comparative study could give a valuable insight into the role of eye movements in glaucoma etiology.

The aim of this study was to map *in vivo* deformation and strain of the ONH tissues in response to changes in gaze positions (abduction and adduction) and to IOP elevation, in both subjects with HTG and NTG. Similar to our previous works^3, 17, 18^, we employed a digital displacement and strain mapping algorithm on spectral domain optical coherence tomography (OCT) images to quantify *in vivo* ONH strains in each subject. We hypothesize that NTG and HTG subjects may have different sensitivities to different biomechanical loads induced by eye movements.

## Methods

Our goal was to quantitively map and compare 3D ONH deformations in NTG and HTG subjects under the following loads – IOP elevation, adduction and abduction. To this end, we first imaged each subject’s ONH in primary gaze using OCT, and subsequently, under each load. ONH tissue deformations were mapped using a digital volume correlation (DVC) algorithm applied to pairs of OCT volumes. Such deformations were then statistically compared across groups (NTG vs HTG), ONH regions, and ONH tissues. Below is a detailed description of our methodology.

### Subjects Recruitment

We recruited 114 subjects with HTG and 114 with NTG from glaucoma clinics at the Singapore National Eye Centre. We included subjects aged more than 50 years old, of Chinese ethnicity (predominant in Singapore), with a refractive error of ±3 diopters, who are currently receiving IOP-lowering medications. We excluded subjects who underwent prior intraocular/orbital/brain surgeries, subjects with history of strabismus, ocular trauma, ocular motor palsies, orbital/brain tumors; with clinically abnormal saccadic or pursuit eye movements; subjects with poor LC visibility in OCT (<50% *en-face* visibility); subjects with known carotid or peripheral vascular disease; or with any other abnormal ophthalmic and neurological conditions. Glaucoma was defined as glaucomatous optic neuropathy, characterized as loss of neuroretinal rim with vertical cup-to-disc ratio >0.7 or focal notching with nerve fiber layer defect attributable to glaucoma and/or asymmetry of cup-to disc ratio between eyes >0.2, with repeatable glaucomatous visual field defects (independent of the IOP value) in at least 1 eye. NTG subjects had low/normal IOP (<21 mmHg) before treatment in the study eye; HTG subjects had elevated IOP (>=21 mmHg) before treatment in the study eye. NTG and HTG categorization was established based on IOP values obtained from Goldmann tonometry.

Each subject underwent the following ocular examinations: (1) measurement of refraction using an autokeratometer (RK-5; Canon, Tokyo, Japan) and (2) measurement of axial length, central corneal thickness and anterior chamber depth using a commercial device (Lenstar LS 900; Haag-Streit AG, Switzerland). For each tested eye we performed a visual field test using a standard achromatic perimetry with the Humphrey Field Analyser (Carl Zeiss Meditec, Dublin, CA).

This study was approved by the SingHealth Centralized Institutional Review Board and adhered to the tenets of the Declaration of Helsinki. Written informed consent was obtained from each subject.

### OCT Imaging

One eye of each subjects was analyzed. If both eyes had similar diagnosis, then we selected the study eye at random for each subject; and the ONH was imaged with spectral-domain OCT (Spectralis; Heidelberg Engineering GmbH, Heidelberg, Germany). The imaging protocol was similar to that from our previous work.^3^ In brief, we conducted a raster scan of the ONH (covering a rectangular region of 15° x 10° centered at the ONH), comprising of 97 serial B-scans, with each B-scan comprising of 384 A-scans (**Figure 1a**). The average distance between B-scans was 35.1 μm and the axial and lateral B-scan pixel resolution were on average 3.87 μm and 11.5 μm respectively. All A-scans were averaged 20 times during acquisition to reduce speckle noise. Each eye was scanned four times under four different conditions – primary OCT position, 20° adduction, 20° abduction and acute IOP elevation. Each subject was administered with 1.0% Tropicamide to dilate the pupils before imaging.

**Figure 1.**
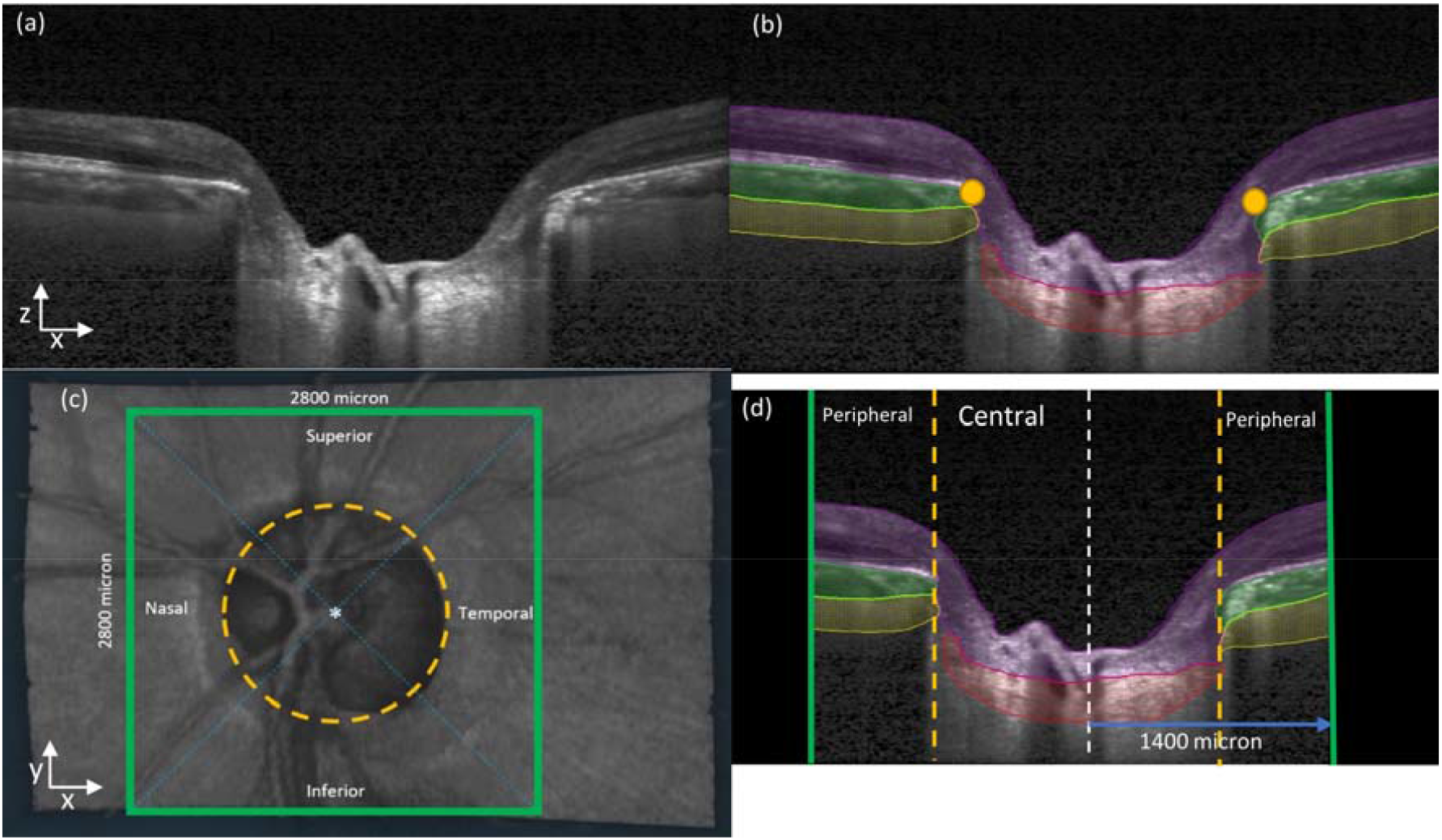
**(a)** A single B-scan obtained from the OCT machine without any image enhancement **(b)** Automatic segmentation of the B-scan in (a). Four tissues were segmented – Pre-Lamina tissue (blue), Choroid (green), Sclera (yellow) and LC (red) In addition, BMOs (orange dots) were automatically marked for each B-scan **(c)** Anterior-surface view (X-Y plane) of the ONH. The ONH center (white star) was identified from the best-fit circle to the BMOs (orange-dotted line). Green square defines our region of interest to be cropped from the OCT volume with 2800μm length on each side. Diagonal blue lines define Superior-Inferior-Nasal-Temporal regions of the ONH. **(d)** A B-scan view (X-Z plane) after we apply cropping to the OCT volumes. Black region was not considered for our deformation tracking. The length from central line (white-dotted line) to the cropping border (green line) is 1400 μm. The area inside the BMO (orange-dotted line) is defined as central region while the area outside the BMO is defined as peripheral region.

### OCT imaging in primary gaze and in Adduction/Abduction positions

In this study, the primary gaze OCT position referred to the eye position during a standard OCT scan. Such a position does not exactly correspond to the primary gaze position as both the pupil and ONH need to be aligned with the OCT objective, inducing a slight eye rotation to the left in a right eye, and vice versa^3^. Amplitudes of horizontal gaze positions reported in this study were therefore with respect to the primary gaze OCT position. Procedures for imaging under different gaze positions have been described in our previous work^3^. Briefly, we employed a custom-built 3D printed rotatable chin rest to induce 20° adduction and 20° abduction and one OCT volume was acquired in each position.

### OCT imaging during acute IOP elevation

For each eye in the primary gaze position, we applied a constant force of 0.65 N to the temporal side of the lower eyelid using an opthalmodynamometer, as per a well-established protocol.^3, 19^ This force raised IOP to about 35 mmHg and was maintained constant throughout the entire OCT acquisition (approximately 3-5 minutes). IOP was then re-assessed with a Tono-Pen (Reichert, Inc.), and the ONH was imaged with OCT in primary gaze position.

### Digital Alignment of OCT volumes

To improve the performance of our deformation mapping protocol, it is first necessary to remove rigid-body translations and rotations that are present due to head and/or eye movements of the subjects in between OCT acquisitions. To this end, each OCT volume under a biomechanical load (adduction, abduction, or elevated IOP) was digitally aligned with its corresponding primary gaze OCT volume using a commercial software Amira (version 2020.1, FEI, Hillsboro, Oregon, USA), as described in our previous publication.^20^

### ONH Reconstruction through Automatic Segmentation

For each ONH, we automatically segmented the following tissue groups - the pre-lamina tissue (PLT, inclusive of retina), the choroid, the sclera and the LC (**Figure 1a-b**) - using a deep-learning algorithm similar to that designed in our previous work.^21, 22^ This was done, so that we can report deformation and strains for each tissue group. Bruch’s membrane opening (BMO) points were also automatically extracted with a custom algorithm. Note that BMO points lie within a plane (the BMO plane), and such a plane can be used as a horizontal reference plane for each ONH.

### In Vivo Displacement and Strain Mapping of the ONH

We used a commercial DVC module (*Amira*. (2020.3). Waltham, Massachusetts: Thermo Fisher Scientific) to map the three-dimensional deformations of the following OCT volume pairs – **(1)** primary gaze vs acute IOP elevation, **(2)** primary gaze vs adduction, and **(3)** primary gaze vs abduction – for each patient. The working principle of this commercial DVC module is similar to our prior DVC implementation^18^, albeit with an improved speed efficiency. Details of the DVC algorithm used in this study is provided in **Appendix A**. Briefly, each ONH morphology was sub-divided into ∼4000 cubic elements, and ∼3,500 nodes (points), at which locations 3D displacements (vectors) were mapped following a change in load (i.e., IOP, adduction, or abduction). We then derived the effective strain from the 3D displacements. The effective strain is a convenient local measure of 3D deformation that takes into account both the compressive and tensile effects. In other words, the higher the compressive or tensile strain, the higher the effective strain. Details of the strain derivation is provided in **Appendix B**, and further validation of the DVC and its effects on strain in **Appendix C**.

### Definition of ONH regions

To ensure un-biased comparisons between groups (NTG vs HTG) for 3D deformations/strains, we first limited our *en-face* field-of-view to a region of 2800 × 2800 μm^2^ centered on the BMO center for all ONHs. Each ONH was further divided into eight regions – inferior, superior, nasal, and temporal from either the central (within the BMO circle), or peripheral (outside the BMO circle) regions (**Figure 1c**). It should be noted that the central region mainly consists of the PLT and LC, whereas the peripheral region of the retina, choroid and sclera (**Figure 1d**).

### Statistics

Statistical analyses were performed using MATLAB (version 2018a, The MathWorks, Inc., Natick, Massachusetts, USA). Similar to our previous work,^19^ strains and displacements were defined as continuous variable and the subjects’ diagnoses (HTG and NTG) were defined as categorical variables. We used independent samples t-test to compare the mean values of effective strain and displacements between the two diagnostic groups. Furthermore, we used Wilcoxon Signed Ranked test to investigate the influence of different loads on the median values of effective strain and displacements. To study the variation of displacements and effective strain between each of the defined ONH regions (**Figure 1c-d**), displacements and effective strain were defined as continuous variable and each region was defined as a categorical variable and the mean and median values of the effective strain were extracted for each of the ONH region. The differences in regional effective strain values were analyzed using generalized estimating equations (GEE), performed using *R* (version 4.0.3; R Foundation, Vienna, Austria) in order to account for inter-region associations. Lastly, to compare the effective strain across different tissues, we also used GEE with effective strain defined as a continuous variable and each tissue type as categorical variables. Statistical significance level for this study was set at *0*.*05*.

## Results

### Demographics and IOP elevation

A total of 228 Chinese subjects were recruited (consisting of 114 subjects with HTG and 114 with NTG). We excluded 10 HTG subjects and 8 NTG subjects from the study due to a low *en-face* LC visibility (<50% of the BMO area) and therefore 104 HTG subjects and 106 NTG subjects were included in the final analysis. Out of 104 HTG subjects, 37 subjects were female. Out of 106 NTG subjects, 49 subjects were female.

There were no significant differences (p>0.05, **Table 1**) across both groups in terms of age [HTG: 69 ± 5, NTG: 67 ± 6], systolic blood pressure [HTG: 141 ± 16 mmHg, NTG: 140 ± 20 mmHg], diastolic blood pressure [HTG: 75 ± 9 mmHg, NTG: 74 ± 9 mmHg], axial length [HTG: 24.2 ± 1.0 mm, NTG: 24.4 ± 1.0 mm], visual field mean deviation [HTG: −7.54 ± 5.05 dB, NTG: −6.56 ± 4.91 dB], pattern standard deviation [HTG: 7.18 ± 3.79 dB, NTG: 7.22 ± 3.05 dB], baseline IOP (on the day of the experiment) [HTG: 17.3 ± 2.9 mmHg, NTG: 16.0 ± 2.5 mmHg], and IOP during ophthalmodynamometer indentation [HTG: 34.5 ± 7.0 mmHg, NTG: 34.8 ± 6.5 mmHg].

**Table 1:** Demographics and clinical characteristics of included study subjects. □IOP values indicated here are measured at the point of the experiment (after glaucoma diagnosis and IOP-lowering treatments for both groups)

| | Mean ± standard deviation or n (%) | | p-value |
| --- | --- | --- | --- |
| Characteristic | HTG | NTG | HTG – NTG |
| Age (year) | 69 ± 5 | 67 ± 6 | 0.10 |
| Sex, female (%) | 32% | 45% | - |
| Systolic blood pressure (mmHg) | 141 ± 16 | 140 ± 20 | 0.91 |
| Diastolic blood pressure (mmHg) | 75 ± 9 | 74 ± 9 | 0.88 |
| Axial Length (mm) | 24.2 ± 1.0 | 24.4 ± 1.0 | 0.13 |
| Visual field, MD (dB) | -7.54 ± 5.05 | -6.56 ± 4.91 | 0.32 |
| Pattern standard deviation (dB) | 7.18 ± 3.79 | 7.22 ± 3.05 | 0.46 |
| Average RNFL thickness (μm) | 59.3 ± 35.5 | 67.4 ± 36.0 | <0.001 |
| Baseline IOP (mmHg)<sup>□</sup> | 17.3 ± 2.9 | 16.0 ± 2.5 | 0.14 |
| IOP (mmHg) with indentation<sup>□</sup> | 34.5 ± 7.0 | 34.8 ± 6.5 | 0.60 |

Retinal nerve fiber layer (RNFL) thickness of the NTG subjects were significantly higher on average as compared to that of the HTG subjects [NTG: 67.4 ± 36.0 μm, HTG: 59.3 ± 35.5 μm, p<0.001].

### IOP induces posterior displacements, while adduction induces anterior displacements in the LC

On average across all subjects, IOP elevation induced posterior displacements of the LC (with respect to the BMO plane) [−5.50 ± 6.73 μm], while abduction and adduction induced anterior displacements [+0.72 ± 9.85 μm and +1.29 ± 6.31 μm respectively] (**Figure 2a**).

**Figure 2.**
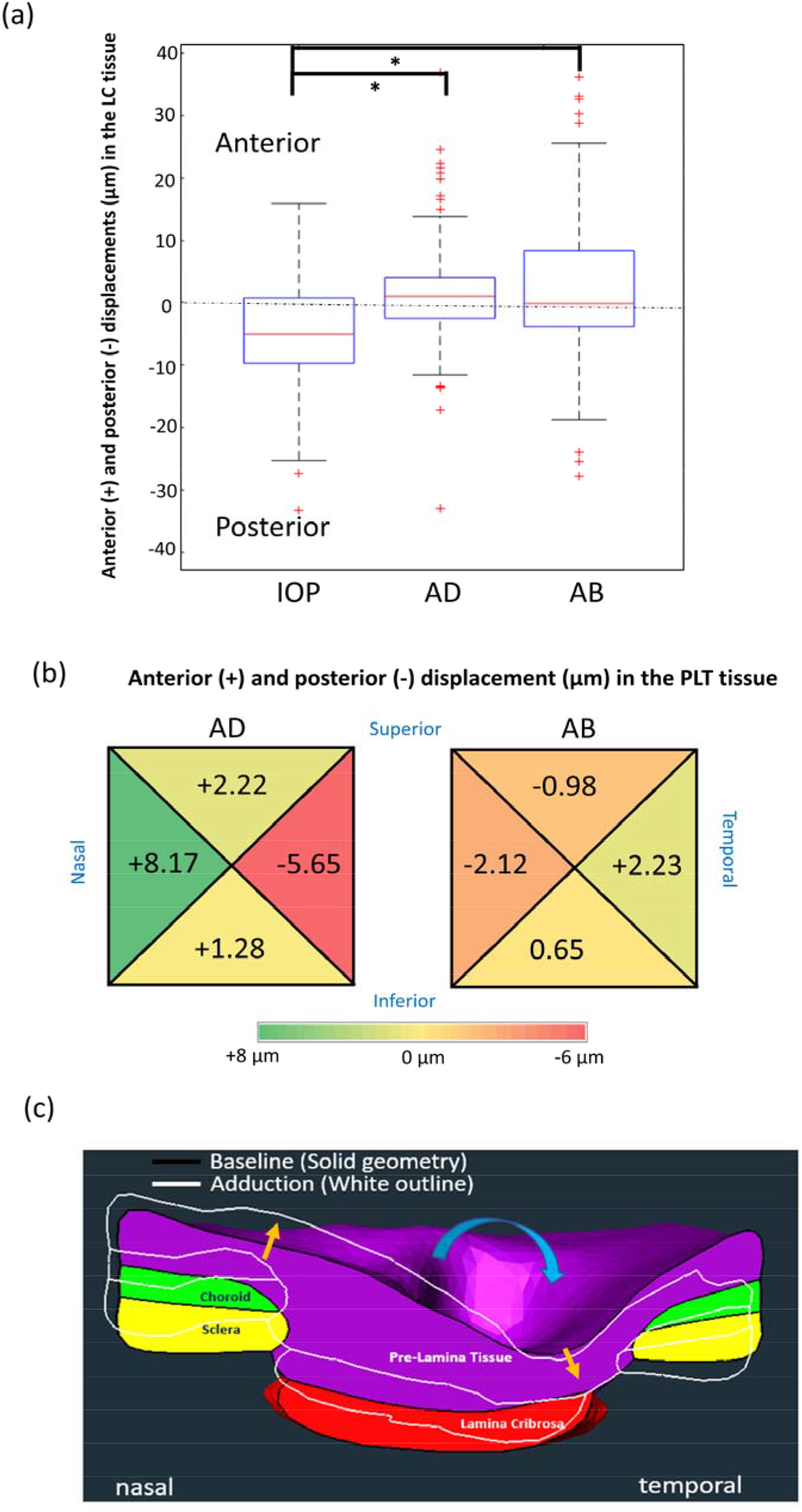
**(a)** A box plot showing anterior-posterior displacement in μm in the LC tissue with respect to each load. (**\*** indicates significant difference at p<0.05) **(b)** A colored-coded plot of regional variations in anterior-posterior displacement (in μm) in PLT with respect to eye positions. **(c)** An example of tissue displacement under Adduction obtained from one subject. Yellow arrows indicate general movement of tissues in nasal and temporal region. Blue arrow indicates globe rotation direction under adduction. AD: adduction, AB: abduction.

### Adduction induces transverse shearing of the PLT

Under adduction, we observed a distinct ‘seesaw’ displacement pattern (i.e., shearing in the transverse plane) in the PLT with, on average, anterior displacements in the nasal region [+8.17 ± 9.31μm] and posterior displacements in the temporal region [−5.65 ± 8.28 μm], with significant difference across two regions (p<0.001) (**Figure 2b-c**). Abduction resulted in an opposite trend of lesser magnitude (posterior displacement nasally [−2.12± 6.53 μm] and anterior temporally [+2.23 ± 6.51 μm]), with significant difference across the two regions (p<0.001). Overall, these trends were also observed for the choroid and sclera.

### IOP and adduction induce equivalently high effective strain in the LC

We observed that both IOP elevation and adduction induce equivalently high effective strain in the LC [4.46 ± 2.4% and 4.42 ± 2.3%, respectively; no significant difference, p = 0.76]. Abduction induced significantly lower effective strain [3.12 ± 1.91%] than both IOP elevation and adduction (p<0.014; **Figure 3**).

**Figure 3.**
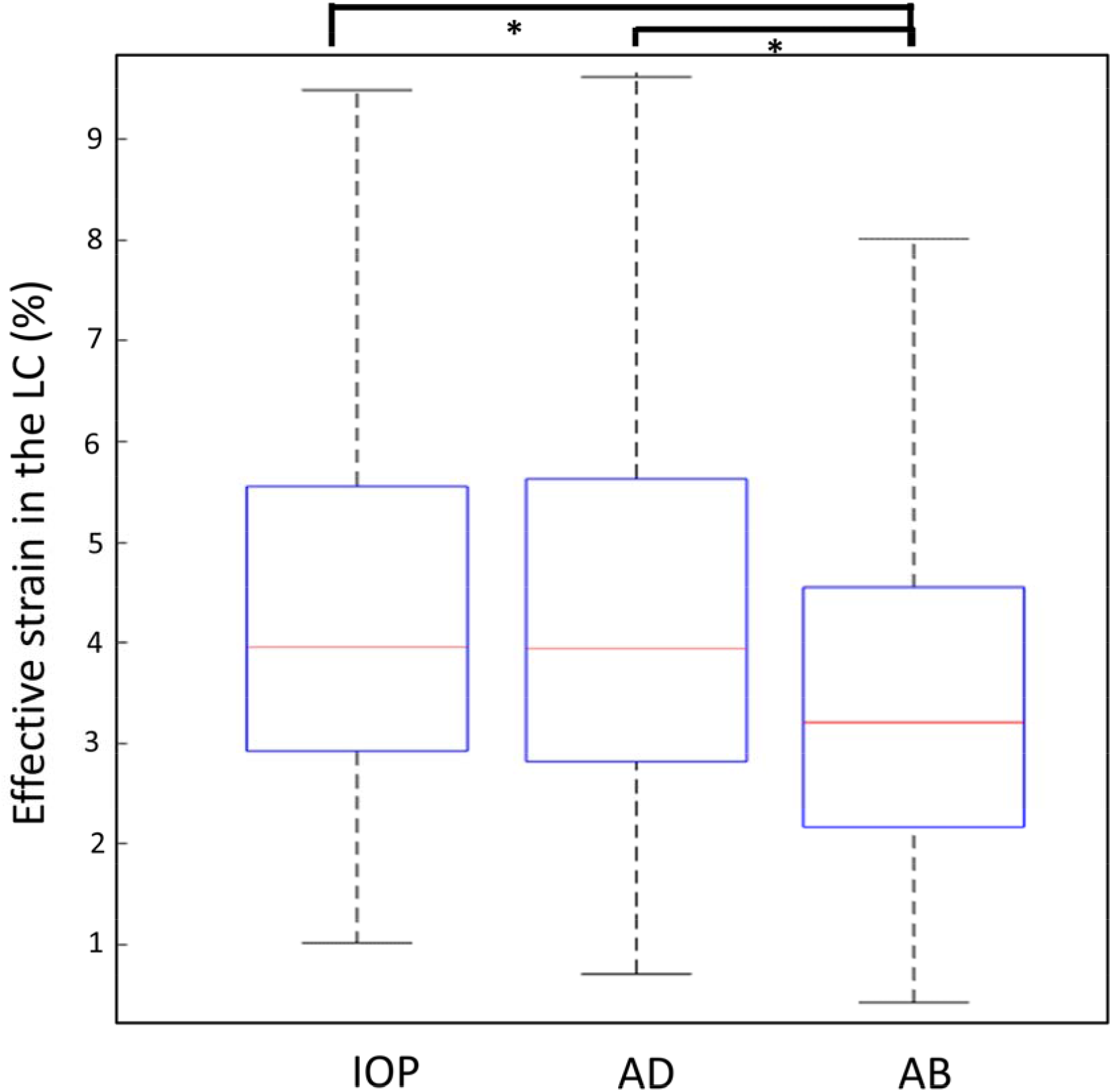
A box plot showing average effective strain in the LC tissue (%) with respect to each load (**\*** indicates significant difference at p<0.05). AD: adduction, AB: abduction.

### NTGs are more sensitive to deformations under adduction, while HTGs under IOP elevation

Under IOP elevation, HTG subjects experienced higher effective strain than NTG in all tissues, with a statistically significant difference observed in the LC tissue [HTG LC: 4.56 ± 1.74% vs NTG LC: 4.12 ± 1.46%, p = 0.047] (**Figure 4a**). Under Adduction, NTG subjects experienced higher effective strain than HTG subjects in all tissues (**Figure 4b**), with a statistically significant difference observed in the LC tissue [NTG LC: 4.93 ± 1.88% vs HTG LC: 4.00 ± 1.40%, p = 0.041] and in the PLT [NTG PLT: 4.56 ± 1.44% vs HTG PLT: 3.86 ± 1.23%, p = 0.002]. Under abduction, no significant differences in effective strain were observed between NTG and HTG subjects.

**Figure 4.**
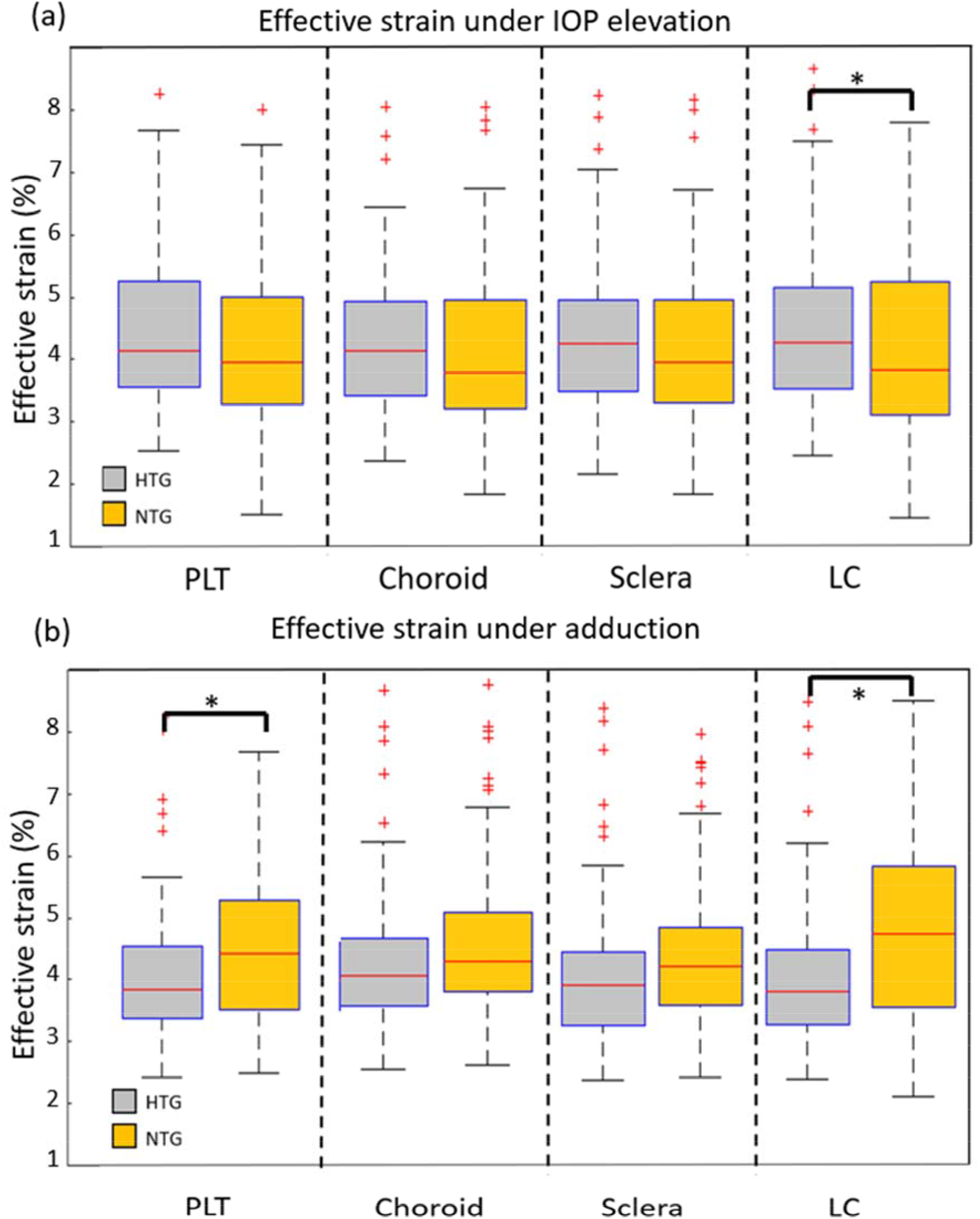
**(a)** A bar chart showing average effective strain in each tissue (under IOP elevation) for each diagnostic group. **(b)** A bar chart showing average effective strain in each tissue (under adduction) for each diagnostic group (**\*** indicates significant difference at p<0.05).

### Regional variations in ONH effective strain

Across all subjects, the central region of the ONH experienced significantly higher effective strain than the peripheral region under IOP elevation [central region: 4.68 ± 1.31% vs peripheral region:3.90 ± 1.13%, p<0.001] and under adduction [central region: 4.53 ± 1.52% vs peripheral region:3.61 ± 1.12%, p<0.001] (**Figure 5**).

**Figure 5.**
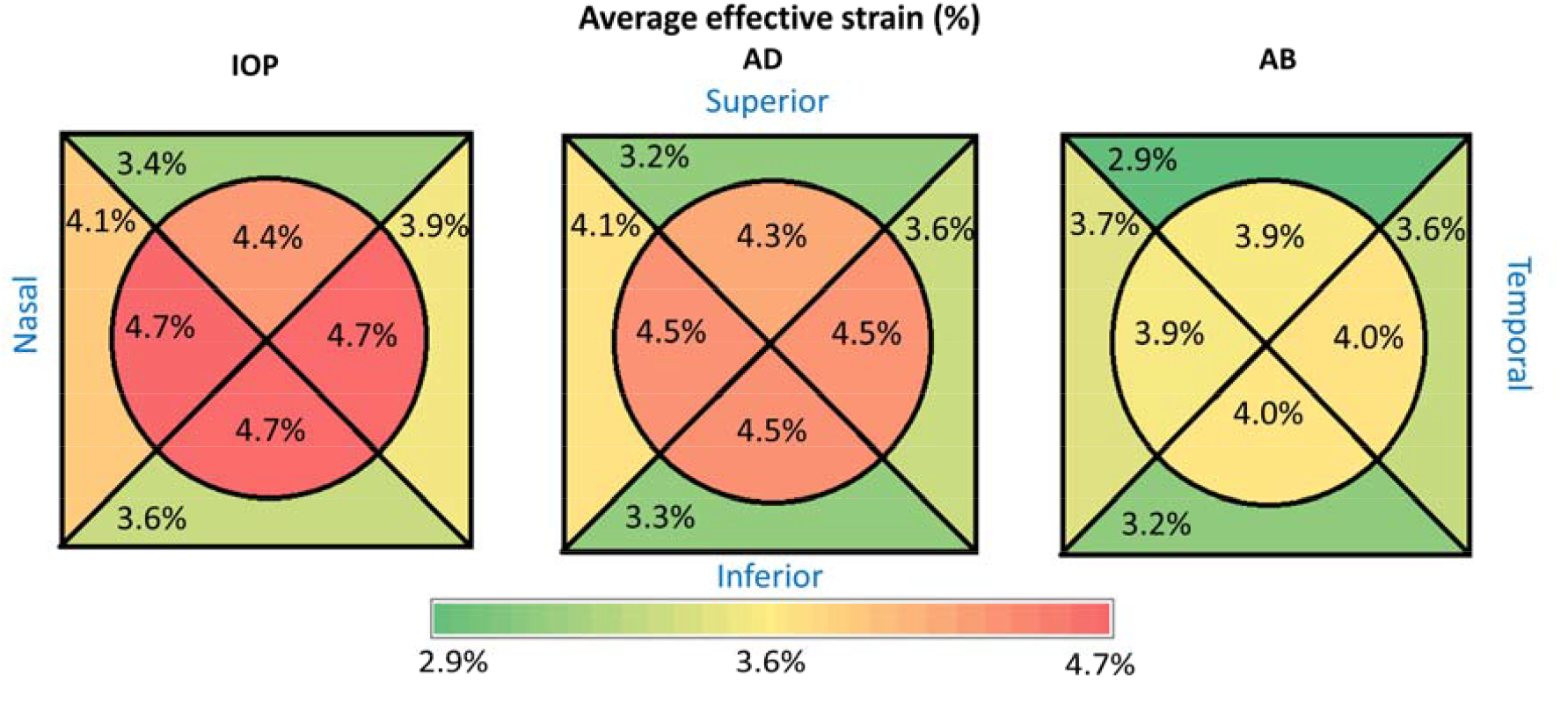
A colored-coded plot of regional variations in average effective strain with respect to each load.

Significantly higher effective strain was observed in the peripheral-nasal region as compared to the peripheral-temporal region under adduction [peripheral-nasal:4.05 ± 1.50% vs peripheral-temporal:3.57 ± 1.20%, p<0.001] (**Figure 5**). For abduction, a homogenous distribution of effective strain was observed with no significant differences across nasal-temporal regions (**Figure 5**).

## Discussion

In this study, we were able to map *in vivo* three-dimensional deformations of the ONH tissues in subjects with HTG and NTG under the presence of biomechanical loads, namely, acute IOP elevation, and optic nerve traction in adduction and abduction. Overall, we found that adduction resulted in large ONH deformations that were on the same order of magnitude as those induced by an acute IOP increase to 35 mmHg. In addition, the ONH of HTG subjects was more mechanically sensitive to IOP compared to whereas that of NTG subjects to adduction. To the best of our knowledge, this is the first study to quantitatively compare *in vivo* ONH deformations between NTG and HTG subjects under different biomechanical loads.

We found that adduction (but not abduction) resulted in ONH deformations and strains that were on the same order of magnitude as those induced by an IOP to 35 mmHg. These findings are consistent with those from our previous studies;^3, 16^ we have simply confirmed this trend in glaucoma eyes. Since adduction stresses the ONH to a similar level as an acute IOP increase, we could potentially consider adduction as a clinical stress test to assess the robustness of the ONH in vivo. The advantage being that imaging the eye in adduction is user-friendly and does not require IOP manipulations that could be of discomfort to the patients. We are currently investigating the relevance of such a stress test in predicting glaucoma progression. We also found that adduction generated significantly higher ONH deformations and strains than abduction; this observation may be explained by the fact that the distance between orbital apex and the ONH is larger following adduction as compared to abduction, which results in a taut optic nerve and a high optic nerve traction during adduction.

Under IOP elevation, we found that the LC of HTG subjects was subjected to significantly higher effective strain than the LC of NTG subjects. In short, HTG subjects were found to be more sensitive to IOP elevation as compared to NTG subjects, and the difference in sensitivity was most pronounced in the LC tissue. Since the LC strains are governed by many factors - for instance, the stiffness of the LC itself^23^, the geometry of the eye^23^, the stiffness of the surrounding sclera^24^ and the complex interactions of the aforementioned parameters - it is difficult to formulate a biomechanical explanation for the observed differences. Experimental and computational modelling studies agree on the importance of sclera being the main eye^24, 25^ load-bearing tissue of the and the factors that determine the load-bearing capacity of the sclera are its shape (thickness and geometry) and its stiffness modulus. A stiffer peripapillary sclera tissue in general was found to reduce the LC.^26, 27^A magnitude of biomechanical strains within the thin posterior sclera was found to deform more than the thick sclera under a given load, resulting in a greater scleral canal expansion and LC deformation.^28^ Indeed, the combinations of other parameters in the ONH could outweigh the contribution of the sclera to the ONH structural strength, sometimes resulting in a high IOP-induced LC strain despite the presence of a stiff sclera. Notwithstanding the complexity of ONH biomechanics, we still believe that the ONH tissues’ stiffnesses and geometries, particularly that of the sclera, are important to formulate a systematic biomechanical model that explains the difference in IOP-induced LC strains between NTG and HTG subjects. To the best of our knowledge, the tissues’ properties of NTG eyes have not been studied. Future work which aims to quantify the tissue properties of NTG eyes may allow us to develop a deeper understanding of the clinically relevant factors that moderate the influence of IOP elevation in NTG subjects.

In this experiment, NTG subjects performing adduction experienced higher effective strain than HTG subjects across all tissues, with a statistically significant difference in the PLT and LC. To explain this, we may again consider the hypothesis that a taut optic nerve acts on the ONH under adduction. It follows then that the degree of force exerted by a taut optic nerve depends on its stiffness (i.e. a stiffer optic nerve will transfer more force to the ONH)^4^ and its length (i.e. a shorter optic nerve will have to stretch more under adduction and will exert more force on ONH). Thus, it is possible that stiffer and/or shorter optic nerves contribute to the greater sensitivity of NTG eyes to adduction-induced deformation. Unfortunately, both the stiffness and length of the optic nerve have not been extensively studied and further studies are warranted.

Interestingly, we observed a transverse shearing of the PLT tissue under adduction, with the ONH tissue in the nasal region displaced anteriorly and the ONH tissue in the temporal region displaced posteriorly (**Figure 2c**). This phenomenon was also observed in abduction (in the opposite direction), although the magnitude of displacement was significantly less than that of adduction. Similar ONH tissue displacement patterns in the nasal-temporal regions under adduction had been documented via a geometric characterization of B-scan images by Wang et al.^3^ and Lee at al.^29^ Demer et al. had also shown in an MRI study that, under adduction, the optic nerve becomes more taut on the temporal side, exerting its ‘pulling’ force mostly on the temporal peripapillary tissue^4, 30^ while indirectly causing the nasal tissue to displace anteriorly. The anterior displacement of the nasal tissue may be explained by the redistribution of cerebrospinal fluid to the nasal side of the ON as the ON sheath on the temporal side is flattened during adduction – this phenomena effectively elevates the CSFP on the nasal side of the ON and the elevated CSFP ‘pushes’ the nasal peripapillary tissue anteriorly.^30^ Our study is the first to show a transverse shearing under adduction in a large number of glaucoma subjects and our observations seem to further support the notion that the optic nerve is acting strongly on the ONH during adduction.

Despite the observation in this study that adduction is significantly influencing the ONH biomechanics, especially for the NTG subjects, more investigations are needed to associate the role of eye movements to the development of glaucoma. Clearly, susceptibility of an eye to adduction alone is not sufficient to cause glaucoma; for instance, a person with a convergent squint (i.e., with one or both eyes in a permanent state of adduction) is not known to be under risk of developing glaucoma. We suspect that if adduction is going to cause long term damage to the ONH tissues, it should come from a repetitive movement pattern that occurs in an eye with prior biotechnical susceptibilities (e.g., shorter and/or stiffer optic nerve) to the ONH strains induced by adduction. This effect could also be cumulative over several years. We can allude this phenomenon to a repetitive neurotrauma that leads to a neurodegenerative disease; for instance, the amyotrophic lateral sclerosis (ALS) occurring in football players^31,32^ that may be caused by repeated head traumas throughout the players’ career. A longitudinal study of glaucoma progression in NTG subjects in relation to the extent of ONH strains under adduction may help to elucidate the causal relationship between eye movements and glaucoma disease.

In terms of demographics and clinical characteristics, we found no significant differences between HTG and NTG subjects, except for the average RNFL thickness (**Table 1**). Even though functional damage or glaucoma severity (as indicated by visual field indexes) were comparable between the two groups, the degree of structural damage (as assessed through RNFL thickness) was different. As glaucoma eyes remodel differently at different stages of damage,^33^ this may have affected our comparisons, and future work that would consider similar structural damage is warranted.

Several limitations in this study warrant further discussion. First, we did not consider (or measure) CSFP in this study. Several studies suggested that NTG patients had lower CSFP as compared to HTG^34–36^ and a low CSFP has become one of the main suspects in the pathogenesis of NTG. In an experimental study, a low CSFP has been shown to increase the translaminar pressure at the LC, leading to a similar glaucomatous optic neuropathy as observed during a development of glaucoma with elevated IOP.^37^ Unfortunately, a non-invasive measurement of CSFP *in vivo* is still not feasible and CSFP is usually estimated from other surrogate measurements, such as an orbital subarachnoid space width^38^ or from a regression model based mainly on values of blood pressures.^39^ If we were able to measure and vary CSFP *in vivo*, we could then investigate the CSFP’s influence on the ONH deformation. We have shown here that the ONH of NTG and HTG subjects deform differently under adduction and such comparison with respect to changes in CSFP would be illuminating to the pathogenesis of NTG. In addition, the differences in CSFP between NTG and HTG may also contribute to the differences in their biomechanical responses when subjected to IOP elevation as observed in this study.

Second, our method of IOP elevation (ophthalmodynamometer) had a considerable degree of uncertainties. Even though we tried to keep the force applied at a constant level, the actual IOP raised still depended on the structural properties of each subject’s eye. This resulted in variations in elevated IOP in both groups of subjects (**Table 1**). To take into account such variations, we then performed an analysis to normalize the strain according to the actual IOP raised in each subject (**Appendix D**; using a linear assumption that could be justified according to Midgett et al.^40^) and found that the average adjusted LC effective strains under IOP elevation were still significantly higher for HTG subjects as compared to NTG subjects (HTG: 6.1±3.2%, NTG: 4.6±2.3%, p = 0.006).

Third, our study was limited to subjects of Chinese ethnicity that were more than 50 years old. Since age is a well-known factor affecting the biomechanical properties of the eyes^33, 41, 42^ and ethnicity is also another important factor that could affect glaucoma pathogenesis^43^, further studies should investigate if the reported differences in biomechanical responses are also present in other demographics.

Fourth, the OCT resolution and signal quality of the posterior portion of the eye (beyond the LC) were poor. Therefore, we were not able to investigate the local strains of the optic nerve in situ, as well as its sheath insertion into the sclera. We suspect that the local strains at the site will be large, and this could contribute to focal defects observed in NTG patients under adduction. In addition, there were differences in terms of scan resolutions across each dimension (i.e., 11.3 micron for the lateral resolution and 3.87 micron for the axial direction). One main implication of the differences in resolution was that the magnitude of displacement error in each direction would be different. For instance, our displacement measurements were approximately 3 times more accurate in the axial direction as compared to the lateral direction. Since our effective strain was an ‘average’ measure across all dimensions, its accuracy would largely be limited by the dimension which had the worst resolution. For instance, an average displacement error (3 micron) in the lateral direction would result in approximately 0.6% error in effective strain, whereas an average displacement error (1 micron) in the axial direction would result in approximately 0.2% error in effective strain.

Fifth, the duration of IOP elevation in our study was short (each patient was subjected to the ophthalmodynamometer procedure for no longer than 5 minutes). This time duration may not be enough to evoke a steady-state deformation response from the tissue. It is likely that we imaged the ONH while it was still in the process of responding to the applied load. However, from our data, we can still clearly observe the influence of IOP elevation on the ONH via a distinct posterior deformation observed in the tissues. Over a long period of time, there may be structural changes in the ONH that may mitigate against such marked deformations. This may explain the different rates of progression observed in glaucoma patients early on and later on in their disease.

Sixth, the displacement and effective strain errors from our DVC method were non-zero. From our validation study, we found that the errors were acceptable when we conducted deformation tracking on repeated primary gaze scans of healthy subjects, with displacement errors of less than 30% of the voxel resolution and the effective strain error of less than 1.2% (**Appendix C**). The errors observed here could arise from various sources such as OCT registration errors (intrinsic to the OCT machine), rotation of subjects’ head during OCT acquisition, OCT speckle noise and IOP fluctuations from ocular pulsations^44^, all of which were difficult to control. Even though our reported baseline strains error of 1.2% was close to the magnitude of differences in average strains between the two groups, we arrived at our main conclusions (i.e., significant differences in strains between HTG and NTG), using a t-test with an appropriate p-value, as opposed to directly comparing the average value of strains. Thus, our main conclusions were valid within the assumptions of t-test (i.e., normal distribution of both population, random sampling of the population etc.).

In conclusion, we measured the *in vivo* deformation of ONH tissue in 114 subjects with HTG and 114 with NTG in response to IOP elevation and horizontal eye movements. We found that **(1)** adduction induced effective strain that was comparable to that induced by IOP, **(2)** NTG subjects experienced significantly higher strains due to adduction compared to HTG subjects and **(3)** HTG subjects (specifically at the LC) experienced significantly higher strain due to IOP elevation compared to NTG subjects. Our results suggest that NTG and HTG subjects have distinct responses to IOP elevation and adduction, supporting the hypothesis that these two have different etiologies of glaucoma damage.

## Financial Support

This work was supported by (1) the Singapore Ministry of Education, Academic Research Funds, Tier 2 (R-397-000-280-112; R-397-000-308-112) (2) Singapore Ministry of Education, Academic Research Funds, Tier 1 (R-397-000-294-114) (3) the National Medical Research Council (Grant NMRC/STAR/0023/2014) and (4) National Natural Science Foundation of China (12002025).

## Financial Disclosures

No financial disclosures.

## Other acknowledgements

None.

## Notes

### Competing Interest Statement

The authors have declared no competing interest.

## References

1. Burgoyne CF, Downs JC, Bellezza AJ, et al. The optic nerve head as a biomechanical structure: a new paradigm for understanding the role of IOP-related stress and strain in the pathophysiology of glaucomatous optic nerve head damage. Progress in retinal and eye research 2005;24(1):39–73.

2. Greene PR. Mechanical considerations in myopia: relative effects of accommodation, convergence, intraocular pressure, and the extraocular muscles. American journal of optometry and physiological optics 1980;57(12):902–14.

3. Wang X, Beotra MR, Tun TA, et al. In vivo 3-dimensional strain mapping confirms large optic nerve head deformations following horizontal eye movements. Investigative ophthalmology & visual science 2016;57(13):5825–33.

4. Demer JL. Optic nerve sheath as a novel mechanical load on the globe in ocular duction. Investigative Ophthalmology & Visual Science 2016;57(4):1826–38.

5. Demer JL, Clark RA, Suh SY, et al. Magnetic resonance imaging of optic nerve traction during adduction in primary open-angle glaucoma with normal intraocular pressure. Investigative ophthalmology & visual science 2017;58(10):4114–25.

6. Wang X, Rumpel H, Baskaran M, et al. Optic nerve tortuosity and globe proptosis in normal and glaucoma subjects. Journal of glaucoma 2019;28(8):691–6.

7. Demer JL, Clark RA, Suh SY, et al. Optic nerve traction during adduction in open angle glaucoma with normal versus elevated intraocular pressure. Current eye research 2020;45(2):199–210.

8. Kim C-s, Seong GJ, Lee N-h, et al. Prevalence of primary open-angle glaucoma in central South Korea: the Namil study. Ophthalmology 2011;118(6):1024–30.

9. Iwase A, Suzuki Y, Araie M, et al. The prevalence of primary open-angle glaucoma in Japanese: the Tajimi Study. Ophthalmology 2004;111(9):1641–8.

10. Yamamoto T, Kitazawa Y. Vascular pathogenesis of normal-tension glaucoma: a possible pathogenetic factor, other than intraocular pressure, of glaucomatous optic neuropathy. Progress in retinal and eye research 1998;17(1):127–43.

11. Mroczkowska S, Benavente-Perez A, Negi A, et al. Primary open-angle glaucoma vs normal-tension glaucoma: the vascular perspective. JAMA ophthalmology 2013;131(1):36–43.

12. Lee SH, Kwak SW, Kang EM, et al. Estimated trans-lamina cribrosa pressure differences in low-teen and high-teen intraocular pressure normal tension glaucoma: the Korean National Health and Nutrition Examination Survey. PloS one 2016;11(2):e0148412.

13. Chen BH, Drucker MD, Louis KM, Richards DW. Progression of normal-tension glaucoma after ventriculoperitoneal shunt to decrease cerebrospinal fluid pressure. Journal of glaucoma 2016;25(1):e50–e2.

14. Park JH, Jun RM, Choi K-R. Significance of corneal biomechanical properties in patients with progressive normal-tension glaucoma. British Journal of Ophthalmology 2015;99(6):746–51.

15. Kim YC, Koo YH, Jung KI, Park CK. Impact of posterior sclera on glaucoma progression in treated myopic normal-tension glaucoma using reconstructed optical coherence tomographic images. Investigative ophthalmology & visual science 2019;60(6):2198–207.

16. Wang X, Rumpel H, Lim WEH, et al. Finite element analysis predicts large optic nerve head strains during horizontal eye movements. Investigative ophthalmology & visual science 2016;57(6):2452–62.

17. Girard MJ, Beotra MR, Chin KS, et al. In vivo 3-dimensional strain mapping of the optic nerve head following intraocular pressure lowering by trabeculectomy. Ophthalmology 2016;123(6):1190–200.

18. Girard MJ, Strouthidis NG, Desjardins A, et al. In vivo optic nerve head biomechanics: performance testing of a three-dimensional tracking algorithm. Journal of The Royal Society Interface 2013;10(87):20130459.

19. Beotra MR, Wang X, Tun TA, et al. In vivo three-dimensional lamina cribrosa strains in healthy, ocular hypertensive, and glaucoma eyes following acute intraocular pressure elevation. Investigative ophthalmology & visual science 2018;59(1):260–72.

20. Maes F, Collignon A, Vandermeulen D, et al. Multimodality image registration by maximization of mutual information. IEEE transactions on Medical Imaging 1997;16(2):187–98.

21. Devalla SK, Renukanand PK, Sreedhar BK, et al. DRUNET: a dilated-residual U-Net deep learning network to segment optic nerve head tissues in optical coherence tomography images. Biomedical optics express 2018;9(7):3244–65.

22. Panda SK, Cheong H, Tun TA, et al. Describing the Structural Phenotype of the Glaucomatous Optic Nerve Head Using Artificial Intelligence. American Journal of Ophthalmology 2021.

23. Sigal IA. Interactions between geometry and mechanical properties on the optic nerve head. Investigative ophthalmology & visual science 2009;50(6):2785–95.

24. Sigal IA, Flanagan JG, Ethier CR. Factors influencing optic nerve head biomechanics. Investigative ophthalmology & visual science 2005;46(11):4189–99.

25. Downs JC, Roberts MD, Burgoyne CF. The mechanical environment of the optic nerve head in glaucoma. Optometry and vision science: official publication of the American Academy of Optometry 2008;85(6):425.

26. Coudrillier B, Campbell IC, Read AT, et al. Effects of peripapillary scleral stiffening on the deformation of the lamina cribrosa. Investigative ophthalmology & visual science 2016;57(6):2666–77.

27. Eilaghi A, Flanagan JG, Simmons CA, Ethier CR. Effects of scleral stiffness properties on optic nerve head biomechanics. Annals of biomedical engineering 2010;38(4):1586–92.

28. Jia X, Yu J, Liao S-H, Duan X-C. Biomechanics of the sclera and effects on intraocular pressure. International journal of ophthalmology 2016;9(12):1824.

29. Lee WJ, Kim YJ, Kim JH, et al. Changes in the optic nerve head induced by horizontal eye movements. PloS one 2018;13(9):e0204069.

30. Chang MY, Shin A, Park J, et al. Deformation of optic nerve head and peripapillary tissues by horizontal duction. American journal of ophthalmology 2017;174:85–94.

31. Abel EL. Football increases the risk for Lou Gehrig’s disease, amyotrophic lateral sclerosis. Perceptual and motor skills 2007;104(3_suppl):1251–4.

32. Chio A, Benzi G, Dossena M, et al. Severely increased risk of amyotrophic lateral sclerosis among Italian professional football players. Brain 2005;128(3):472–6.

33. Downs JC. Optic nerve head biomechanics in aging and disease. Experimental eye research 2015;133:19–29. Beijing

34. Wang N, Xie X, Yang D, et al. Orbital cerebrospinal fluid space in glaucoma: the intracranial and intraocular pressure (iCOP) study. Ophthalmology 2012;119(10):2065–73. e1.

35. Jonas JB, Yang D, Wang N. Intracranial pressure and glaucoma. Journal of glaucoma 2013;22:S13–S4.

36. Cha S, Jeong S, Oh H, et al. Estimated cerebrospinal fluid pressure in normal tension glaucoma versus high tension glaucoma in Korean population based study. Investigative Ophthalmology & Visual Science 2015;56(7):5006–.

37. Yablonski M, Ritch R, Pokorny K. Effect of decreased intracranial-pressure on optic disk. Investigative Ophthalmology & Visual Science: LIPPINCOTT-RAVEN PUBL 227 EAST WASHINGTON SQ, PHILADELPHIA, PA 19106, 1979.

38. Xie X, Chen W, Li Z, et al. Noninvasive evaluation of cerebrospinal fluid pressure in ocular hypertension: a preliminary study. Acta ophthalmologica 2018;96(5):e570–e6.

39. Fleischman D, Bicket AK, Stinnett SS, et al. Analysis of cerebrospinal fluid pressure estimation using formulae derived from clinical data. Investigative Ophthalmology & Visual Science 2016;57(13):5625–30.

40. Midgett DE, Quigley HA, Nguyen TD. In vivo characterization of the deformation of the human optic nerve head using optical coherence tomography and digital volume correlation. Acta biomaterialia 2019;96:385–99.

41. Kotecha A. What biomechanical properties of the cornea are relevant for the clinician? Survey of ophthalmology 2007;52(6):S109–S14.

42. Strouthidis NG, Girard MJ. Altering the way the optic nerve head responds to intraocular pressure—a potential approach to glaucoma therapy. Current opinion in pharmacology 2013;13(1):83–9.

43. Sample PA, Girkin CA, Zangwill LM, et al. The african descent and glaucoma evaluation study (ADAGES): Design and baseline data. Archives of ophthalmology 2009;127(9):1136–45.

44. Pitchon E, Leonardi M, Renaud P, et al. First in vivo human measure of the intraocular pressure fluctuation and ocular pulsation by a wireless soft contact lens sensor. Investigative Ophthalmology & Visual Science 2008;49(13):687–.

