## Supplemental Appendix for "Adduction Induces Large Optic Nerve Head Deformations in Subjects with Normal Tension Glaucoma"

A. **Details of DVC algorithm**

We used a commercial software module (Amira *XDigitalVolumeCorrelation* Extension) to map the three-dimensional deformation of the following OCT volume pairs – (1) baseline OCT vs acute IOP elevation (2) baseline OCT vs adduction, and (3) baseline OCT vs abduction – for each patient. The software utilized a robust Finite Element (FE)-based DVC approach which is often called a ‘global’ approach.^24, 25^ Unlike the ‘local’ DVC approach used in our previous works,^26^ a FE-based DVC requires the FE mesh of the entire region of interest as a method of discretization, in contrast to a ‘sub-volumes’ region of interest for the local approach. Therefore, in the global approach, the whole volume of interest is considered in each iteration and the method used by Amira *XDigitalVolumeCorrelation* was described in detail by Hild et al.^24^ Briefly, FE-based DVC problem consisted of solving the following equation –

$$f\left( x \right)=g\left( x+u\left( x \right) \right) Eq. 1$$

where $x$ is a position of any voxel, $f$ is reference volume voxel intensity (in grayscale), $g$ is deformed volume voxel intensity (in grayscale) and $u$ is the unknown displacement field. Solving the above equation is equivalent to minimizing the sum of squared differences of the correlation residual $\Phi_{c}(x)$ –

$$\Phi_{c}\left( x \right)=|f\left( x \right)-g\left( x+u\left( x \right) \right)| Eq. 2$$

A weak formulation based on C8 finite element (and its associated trilinear shape function) was used to discretize the problem^25^ and to generate the following linear system to be solved –

$$\left[ M \right]\left\{ \delta u \right\}=\left\{ b \right\}Eq. 3$$

where $\left[ M \right]$ is the DVC matrix, $\{\delta u\}$ is the correction vector to vector $u$ at each iteration and $\{b\}$ is the DVC vector that needs to be minimized.

The algorithm converges when $\delta u$ is smaller than a certain tolerance. To summarize, *XDigitalVolumeCorrelation* needs three main inputs – (1) hexahedron mesh of the ROI (2) convergence criterion which is the tolerance ($\delta u$), and (3) a baseline image volume and a deformed image volume. In this study, we generated a hexahedron mesh using AMIRA software mesh generator by specifying a mesh size of 100x100x100 microns or approximately 9x3x26 voxels in X, Y and Z directions for each volume. This resulted in a hexahedron mesh with approximately 3500 nodes for an average volume (**Figure A1a-b**). The DVC algorithm calculated the displacements of each cube’s vertices (the cube’s corners). We set the tolerance of $\delta u$ to be 1x10^-4^ for each volume.

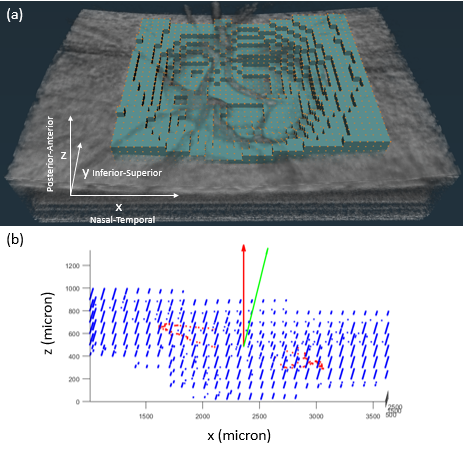

**Figure A1 (a)** 3D volumetric view of OCT scans (in grayscale). Orange dots are nodes (approximately 3500 nodes for each volume) of the hexahedron tracking mesh. Teal boxes represent the elements of the hexahedron tracking mesh. The dimension of each element is 100x100x100 µm in X, Y and Z directions. **(b)** Prior to strain calculation, the nodes (in blue) and their respective displacements are re-oriented with respect to BMO (red dots). Green vector represents the normal vector of a plane fitted to BMO coordinates. Red vector represents the normal vector of X-Y plane. We performed 3D rotation to the entire volume to rotate the green vector to the red vector.

B. **Derivation of strains**

From the displacement field, we calculated a deformation tensor (*F*) and we derived a Green Lagrange strain tensor consisting of 6 components (Exx, Eyy, Ezz, Exy, Eyz, Exz) according to the following equation –

$$E = \frac{1}{2}\left( transpose\left( F \right)\times F-I \right)$$

where E is Green Lagrange strain tensor and I is identity matrix. In this study, we reported effective strain as the main measure of strain within a tissue node. We calculated effective strain from the principal components (E_1_, E_2_, E_3_) of the Green Lagrange strain tensor (E) according to the following equation –

$$E_{eff} =\sqrt{\frac{{(E_{1}-E_{2})}^{2}+{(E_{1}-E_{3})}^{2}+{(E_{2}-E_{3})}^{2}}{2}}$$

where $E_{eff}$ is the effective strain. Note that effective strain is a single positive index that represents both compressive and tensile effects. In other words, higher compressive strain or tensile strain will result in a higher effective strain.

C. **Validation of DVC method**

The DVC algorithm baseline displacements and strains error were estimated from the following artificial cases (performed on a baseline volume of healthy subjects) – (1) 20-micron rigid body translation along a positive x-direction, (2) a 2-degree clockwise rotation about the center of the ONH, (3) 4% tension, (4) 4% compression along the x-direction, (5) radial expansion from the geometric center of the LC with a mean effective strain of approximately 4%, (6) 3% compression along the z-direction, (7) a combination of case 5 and 6 and (8) case 6 with an addition of gaussian noise (mean 0 and standard deviation of 5%). We also estimated the baseline error due to variation in subject’s body position and motion between each scan by comparing repeated baselines (N = 3) scans from a single subject. We quantify the errors in terms of the following parameters: displacement magnitudes in the X, Y and Z direction and effective strains.

For case (1), the average error in displacements were 0.086 ± 0.10 micron in the x-direction, 0.028 ± 0.27 micron in the y-direction, 0.035 ± 0.33 micron in the z-direction and the average error in effective strain value was 0.0024 ± 0.001%. For case (2), the average error in displacements were 0.024 ± 0.66 micron in the x-direction, 0.033 ± 0.43 micron in the y-direction, 0.017 ± 0.068 micron in the z-direction and the average error in effective strain value was 0.0070 ± 0.005%. For case (3), the average error in displacements were 0.26 ± 0.65 micron in the x-direction, 0.002 ± 0.016 micron in the y-direction, 0.015 ± 0.19 micron in the z-direction and the average error in effective strain value was 0.0003 ± 0.002%. For case (4), the average error in displacements were 0.36 ± 0.71 micron in the x-direction, 0.031 ± 0.019 micron in the y-direction, 0.011 ± 0.22 micron in the z-direction and the average errors in effective strain value was 0.0025 ± 0.002%. %. For case (5), the average error in displacements were 0.44 ± 0.76 micron in the x-direction, 0.042 ± 0.033 micron in the y-direction, 0.018 ± 0.31 micron in the z-direction and the average errors in effective strain value was 0.0065 ± 0.003%. %. For case (6), the average error in displacements were 0.015 ± 0.22 micron in the x-direction, 0.011 ± 0.018 micron in the y-direction, 0.033± 0.35 micron in the z-direction and the average errors in effective strain value was 0.007 ± 0.003%. %. For case (7), the average error in displacements were 0.51 ± 0.31 micron in the x-direction, 0.032 ± 0.018 micron in the y-direction, 0.035± 0.35 micron in the z-direction and the average errors in effective strain value was 0.0074 ± 0.003%. For case (8), the average error in displacements were 0.65 ± 0.37 micron in the x-direction, 0.035 ± 0.022 micron in the y-direction, 0.038± 0.37 micron in the z-direction and the average errors in effective strain value was 0.0077 ± 0.003%.

Overall, for the artificial deformation cases, the maximum error in displacements along the X, Y and Z direction was less than 5% of the voxel resolution. The maximum error in effective strain was 0.5%.

For three repeated baseline scans of a healthy subject (N1, N2, N3), the average displacement errors for the pair N1-N2 were 3.2 ± 3.8 micron in the x-direction, 3.3 ± 5.0 micron in the y-direction, 1.2 ± 1.9 micron in the z-direction and the average error in effective strain was 1.0 ± 0.1%. For the pair N1-N3, the average displacement errors were 1.9± 3.1 micron in the x-direction, 5.3 ± 4.8 micron in the y-direction, 1.5 ± 2.8 micron in the z-direction and the average error in effective strain was 0.9 ± 0.08%. For the pair N2-N3, the average displacement errors were 3.4± 3.2 micron in the x-direction, 5.5 ± 4.3 micron in the y-direction, 1.5 ± 2.9 micron in the z-direction and the average error in effective strain was 1.1 ± 0.08%. Overall, for the repeated baseline scans of a single subject, the maximum error in displacements was approximately 30% of the voxel resolution and the maximum error in effective strain was 1.2%.

D. **Adjusting Strains to the IOP-raised**

Since we measured the level of raised IOP, we agree that we should incorporate this information into our analysis. To the best of our knowledge, there is currently no mathematical formulation that represents the relationship between the two parameters (loads induced by IOP elevation and the strains experienced by the ONH). A study by Midgett et al, where the IOP change from baseline (ΔIOP) was varied within [0, 20 mmHg] and the strains in the anterior lamina cribrosa (ALC) was determined using DVC, showed that the relationship between ΔIOP and the strains in the ALC could be adequately characterized by a linear function (p<0.05). In comparison, our range of ΔIOP with the application of ODM was [3, 33 mmHg], which is comparable to the aforementioned study.

Since there are no concrete evidences to suggest a need to use a complex (non-linear) function to relate the two parameters (IOP and LC strains) and by Occam’s razor (the simplest explanation is usually the best one), we assumed that a linear approximation could first be used for LC strains in our range of elevated IOP [18 mmHg, 53 mmHg].

Accordingly, we normalized the strains across each subject (to account for the variations in elevated IOP) based on the average value of the elevated IOP (35 mmHg) according to the following equation -

$$Adjusted Effective Strains=Measured Effective Strain \times\frac{35 mmHg}{Measured elevated IOP}$$

We found that under IOP elevation, the average adjusted LC effective strains were 6.1±3.2% for HTG subjects and 4.6±2.3% for NTG subjects and that there were statistically significant differences between the two groups (p = 0.006).

In other words, our conclusions remain the same even after adjusting for the IOP increase effect.
